# Inhibition mechanism of pancreatic KATP channels by centipede toxins

**DOI:** 10.1101/2025.10.27.684718

**Authors:** Tianyi Hou, Mengmeng Wang, Yongmei Tan, Chengpeng Fan, Lei Chen

## Abstract

The pancreatic ATP-sensitive potassium (KATP) channel acts as a crucial metabolic sensor by regulating insulin secretion to maintain whole-body energy homeostasis. Gain-of-functional mutations in this channel lead to neonatal diabetes mellitus, a rare disorder in which certain mutants demonstrate resistance to standard sulfonylurea therapy. Recent studies have identified a centipede toxin, SpTx1, as a potent inhibitor of both human pancreatic KATP channels and their gain-of-functional mutants. This toxin stimulates insulin secretion, offering a promising therapeutic strategy for neonatal diabetes. Nevertheless, the molecular mechanism by which SpTx1 inhibits the KATP channel has remained elusive. Here, we report the crystal structure of SpTx1 and the cryo-electron microscopy (cryo-EM) structure of the KATP channel in complex with SpTx1. Our results demonstrate that SpTx1 binds to the extracellular surface of the Kir6.2 pore-forming subunit of the KATP channel. Multiple interactions at the SpTx1-Kir6.2 interface underpin the high affinity and specificity of SpTx1 for human Kir6.2. We show that SpTx1 inhibits potassium currents by physically blocking the ion conduction pore. Furthermore, a structure-guided search identified another centipede toxin, Sm3a, as a novel KATP channel blocker. Together, these findings provide key insights into the inhibitory mechanism of centipede toxins against the human Kir6.2-containing KATP channel and establish a foundation for developing these toxins into potential therapeutics targeting KATP channel-related diseases.

## Main

The activities of KATP channels are inhibited by intracellular ATP and activated by Mg-ADP, thereby linking cellular metabolic status to the permeability of potassium ions on the plasma membrane ^1^. As such, KATP channels adjust the cellular membrane potential according to the changes of the intracellular ATP/ADP ratio ^1^. KATP channels are hetero-octamers composed of four central Kir6 pore subunits (either Kir6.1 or Kir6.2) and four regulatory SUR subunits (SUR1 or SUR2) ^1^. Specifically, pancreatic KATP channels are formed by Kir6.2 subunit and SUR1 subunit and play a critical role in regulating insulin release^1–3^. When blood glucose levels are elevated, the cellular metabolism in pancreatic β cells is enhanced, and intracellular ATP concentration is increased, which inhibits the KATP channel^1^. This inhibition results in plasma membrane depolarization and the activation of voltage-gated calcium channels, leading to calcium influx and insulin release^1^. Therefore, KATP inhibitors such as sulfonylureas and glinides are commonly used as insulin secretagogues to block KATP channel activity and enhance insulin secretion^4^.

Gain-of-function mutations in pancreatic KATP channels result in reduced ATP sensitivity and impaired insulin release in response to increased blood glucose, leading to neonatal diabetes in humans^5^. More than 80 gain-of-function mutations in the Kir6.2 channel has been found to lead to neonatal diabetes, exhibiting varying degrees of disease severity ^5^. These range from transient neonatal diabetes mellitus (TNDM) and permanent neonatal diabetes mellitus (PNDM) to severe developmental delay, epilepsy, and neonatal diabetes (DEND) syndrome ^5^. While some patients with neonatal diabetes can achieve adequate blood glucose control with sulfonylurea therapy, others, particularly those with mutations such as L164P, C166Y, I296L, and G334D, exhibit resistance to this treatment^5^.

Structural studies have shown that sulfonylureas and glinides bind to the central pocket of SUR1 in the transmembrane domain, exerting their inhibitory effects on the activity of Kir6.2 allosterically^4,6–12^. Mechanistically, they not only block the dimerization and activation of the nucleotide-binding domains (NBDs) of SUR1 by Mg-ADP but also recruit the N-terminal peptide of Kir6.2 (KNtp) to allosterically close the pore^4,7,10^. Mutations on SUR1 or Kir6.2 that severely disrupt these two mechanisms abolish the inhibitory function of sulfonylureas, rendering patients insensitive to this therapy^5^. For these sulfonylurea-resistant mutations, a promising alternative strategy to inhibit KATP channels is to directly block the Kir6.2 pore.

It has recently been discovered that centipede toxin SpTx1 can specifically inhibit human Kir6.2 (*hs*Kir6.2) with high specificity and potency^13^. Moreover, SpTx1 also blocks Kir6.2 mutants associated with neonatal diabetes that are resistant to sulfonylurea therapy^13^. Further in vivo studies have demonstrated that SpTx1 acts as a potent secondary insulin secretagogue, effectively reducing elevated blood glucose levels in diabetic mice, but it has no effect on nondiabetic mice^14^, suggesting SpTx1 as a promising reagent to treat diabetes, especially the sulfonylurea-resistant neonatal diabetes. Despite these progresses, the molecular details of how SpTx1 specifically binds to and inhibits human Kir6.2 remain unknown. Here, we present the crystal structure of SpTx1 and the cryo-EM structure of the SpTx1-KATP channel complex, providing insights into the inhibitory mechanism of SpTx1.

## Results

### Crystal structure of SpTx1

We expressed the SpTx1 toxin recombinantly in *E. coli* and purified it to apparent homogeneity (Figure S1A). Whole-cell patch clamp experiments demonstrated that purified SpTx1 can inhibit currents of the KATP channel formed by human *hs*Kir6.2 and SUR1 from *Mesocricetus auratus* (*ma*SUR1), indicating that the purified toxin is functional (Figure 1A). We then crystallized SpTx1 and determined its crystal structure via X-ray crystallography at a resolution of 1.9 Å (Figure 1B-D and Table S1). In one asymmetric unit, there are six SpTx1 monomers packed in a head-to-tail fashion, forming a hexametric ring (Figure S1B). The structures of each monomer are highly similar (Figure S1C-D). SpTx1 toxin exhibits a spearhead-shaped structure with an α-helix packing against three antiparallel β sheets (Figure 1B). Both the N-terminus and C-terminus of SpTx1 are located on the same side, at the tail of the spearhead (Figure 1B). On the opposite side, a positively charged K15 is located at the tip of the spearhead (Figure 1B). Two disulfide bonds are formed between C22 on α1 and C48 on β3, and between C26 on α1 and C50 on β3, further stabilizing the overall compact structure of SpTx1 (Figure 1B). These two disulfide bonds were absolutely conserved in other centipede toxins that show inhibitory effect against *hs*Kir6.2 ^15,16^ (Figure 1E). Notably, the surface of SpTx1 is highly hydrophilic and rich in positively charged patches (Figure 1C, D), suggesting that polar interactions might play a crucial role in the interactions between SpTx1 and *hs*Kir6.2. Notably, the overall structure of SpTx1 and the position of the Lys at the tip are also similar to another centipede toxin, SsTx from *Scolopendra subspinipes mutilans*, which specifically inhibits KCNQ channel ^17^(Figure S1E).

**Figure 1.**
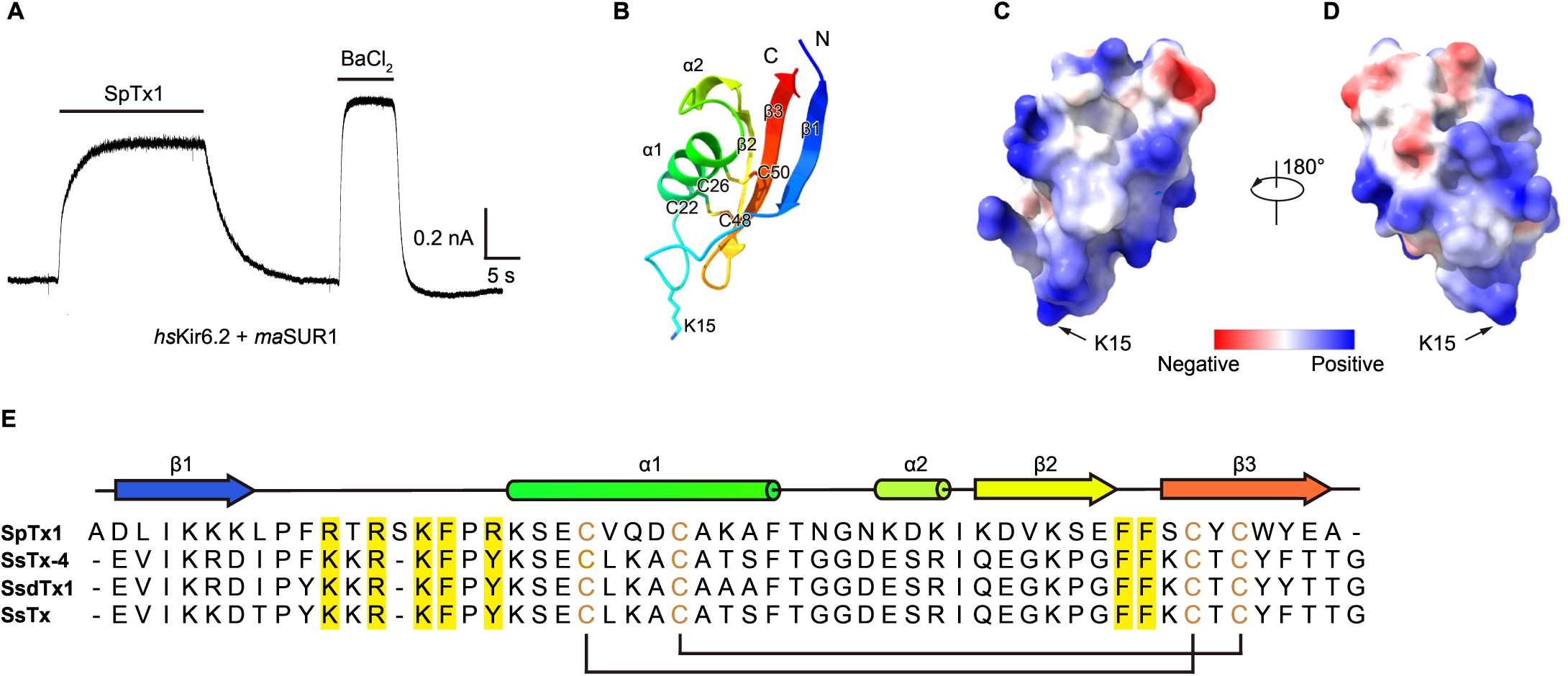
Crystal structure of SpTx1 toxin. (A) 1 µM SpTx1 inhibits the whole-cell current of the *hs*Kir6.2+*ma*SUR1 KATP channel. Cells were clamped at -40 mV. 5 mM BaCl₂ was used to determine the baseline. (B) Crystal structure of SpTx1, colored in rainbow. Two disulfide bonds are shown as sticks. (C) Surface electrostatic potential distribution of SpTx1, shown in red (negative) and blue (positive). SpTx1 is in the same orientation as in (B). (D) A 180°-rotated view of (C). (E) Sequence alignment of SpTx1 and three related toxins: SsTx-4, SsdTx1 (GenBank KC145039), and SsTx (PDB ID: 5X0S). α-helices and β-sheets are represented as cylinders and arrows, respectively, and are colored according to (B). Cysteines are colored in brown, and disulfide bonds are indicated by black lines.

### Structure determination of pancreatic KATP channel in complex with SpTx1

To understand how SpTx1 binds the KATP channel, we sought to determine the cryo-EM structure of their complex. To express the KATP channel for structural studies with SpTx1, we fused *hs*Kir6.2 to the C-terminus of *ma*SUR1 and incorporated an PreScission Protease cleavage site upstream of *hs*Kir6.2 (Figure S2A). This design ensured the 4:4 stoichiometry between *hs*Kir6.2 and *ma*SUR1 and facilitated the correct assembly of the channel ^7,8^. Additionally, cleavage of the *ma*SUR1-*hs*Kir6.2 fusion protein by PreScission Protease released the Kir6.2 N-terminal peptide (KNtp), which is crucial for conferring sensitivity to insulin secretagogues such as glibenclamide^7,8^. We also introduced mutations Q52E in *hs*Kir6.2 and E203K in *ma*SUR1 to enhance the ATP sensitivity of the KATP channel^18^ (Figure S2A). The resulting KATP fusion construct retains the sensitivity to SpTx1 inhibition (Figure S2B), the same as wild type channel (Figure 1A). Following Precission Protease cleavage of the expressed KATP fusion protein and fractionation on size-exclusion chromatography (Figure S2C), we supplemented the protein with purified SpTx1 toxin for cryo-EM sample preparation. We also supplemented the channel protein with ATP and glibenclamide to stabilize the channel in the ATP-bound and glibenclamide-bound closed conformation to obtain a high-resolution structure (Figure S3).

During the cryo-EM image analysis, consensus refinement using C4 symmetry yielded a map with an overall resolution of 2.74 Å (Figure S3); however, the SUR1 ABC transporter region (SUR1_ABC_) appeared blurry, confirming its dynamic interface with the central Kir6.2 and SUR1 TMD0 region (KATP_CORE_) ^12^. Additional focused refinement on SUR_ABC_ with C1 symmetry improved the resolution to 2.87 Å (Figure S3). In the map of SUR_ABC_, we observed the density of glibenclamide in the transmembrane domain, ATP on NBD1, and KNtp in the central cavity of *ma*SUR1 (Figure S4). In the map of KATP_CORE_, we identified toxin density at the extracellular mouth of the Kir6.2 channel (Figure S3), which was absent in previous KATP maps with *hs*Kir6.2 ^19,20^. However, due to the C4 averaging, the density of SpTx1 remained uninterpretable. Further symmetry expansion and focused classification with C1 symmetry on SpTx1 resulted in two distinct 3D classes (Figure S3): one with no SpTx1 bound and another exhibiting prominent SpTx1 density. The class with SpTx1 bound was refined to a resolution of 2.88 Å with C1 symmetry and was subsequently merged with the SUR1_ABC_ maps to generate the composite map for model building and interpretation (Figures 2, S3, S4, and Table S2). In the map, we observed all of the ligands added during sample preparation, including ATP bound in the Kir6.2 subunit, ATP bound in the SUR1 subunit (Figure 2A), and glibenclamide bound in the SUR1 subunit (Figure 2B). We also observed discontinuous density of KNtp in the central cavity of SUR1 (Figure S4B-C), in agreement with its dynamic nature.

**Figure 2.**
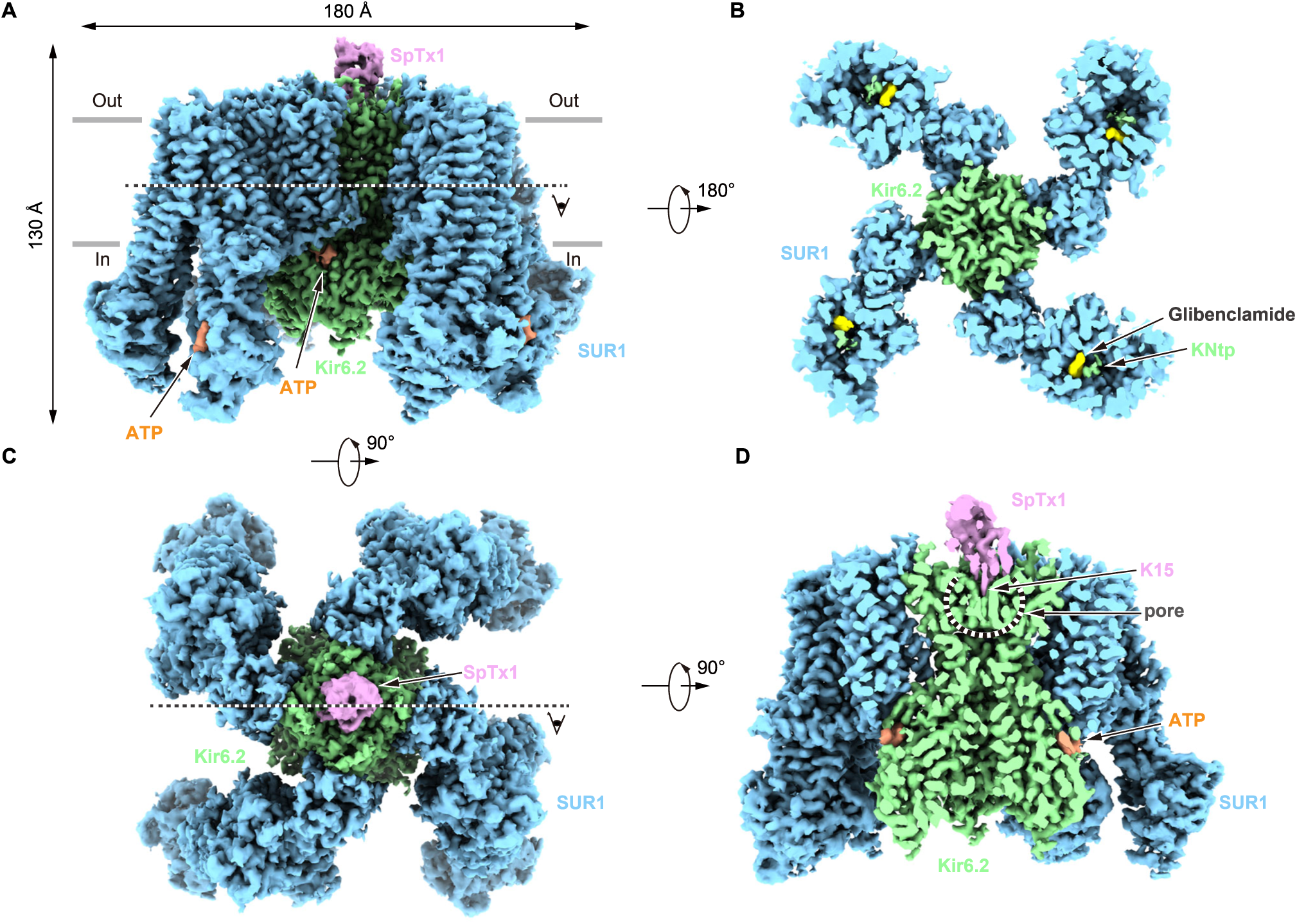
Cryo-EM structure of KATP in complex with SpTx1. (A) Cryo-EM density map of KATP in complex with SpTx1, viewed from the side. The *hs*Kir6.2 subunits, *ma*SUR1 subunits, ATP, and SpTx1 are colored in green, blue, orange, and pink, respectively. (B) A bottom cut-open view, the position is indicated by the dashed line in (A), glibenclamide and KNtp are colored in yellow and green, respectively. (C) Cryo-EM density map of KATP in complex with SpTx1, viewed from the top. (D) A cut-open view from the side at the position of the dashed line indicated in (C), illustrating the binding of SpTx1 at the *hs*Kir6.2 pore.

### Interactions between SpTx1 and *hs*Kir6.2

The structure reveals that SpTx1 binds to the highly negatively charged surface on the extracellular side of the KATP channel (Figure 3A-C and S4). SpTx1 positions its positively charged K15 at its tip into the extracellular mouth of the *hs*Kir6.2 pore (Figure 3B-D and Video S1). The primary amine group of K15 is situated on the four-fold symmetry axis and near the P1 site of the pore of *hs*Kir6.2, where it interacts with the four backbone carbonyl groups of F133 located at the selectivity filter of *hs*Kir6.2 (Figure 3E and S4E). Surrounding K15 are three phenylalanines—F16 on the β1-α1 loop, and F45 and F46 on the β2-β3 loop of SpTx1—which engage with the carbonyl groups of G134 at the selectivity filter of *hs*Kir6.2 via the edges of their benzene rings, and also form hydrophobic interactions with residues at the pore (Figure 3E and S4F). Particularly, both F16 and F45 form hydrophobic interactions with M137 of two adjacent *hs*Kir6.2 subunits (Figure 3E and S4G). Additionally, R11 on the β1-α1 loop of SpTx1 forms one hydrogen bond with the carbonyl group of G135 at the mouth of the Kir6.2 pore (Figure 3F and S4H). Meanwhile, R18 on the β1-α1 loop of SpTx1 interacts with E108 on the S2-S3 turret of one *hs*Kir6.2 subunit (Figure 3G and S4I), and R13 on the same loop further interacts with both E108 and the carbonyl group of G105 on the turret between inner helix (IH) and outer helix (OH) of the adjacent *hs*Kir6.2 subunit (Figure 3F and S4J). To investigate the functional significance of these hydrophilic interactions, we mutated the relevant charged residues to alanines and measured the inhibitory effects of mutant proteins at 1 µM concentration (Figure S5). We found that the K15A mutant of SpTx1 exhibited a great loss of inhibition of the KATP channel (Figure 3H). Similarly, the R11A, R13A, and R18A mutations in SpTx1 all reduced its inhibitory effect to some extent (Figure 3H). Additionally, the E108A mutation in *hs*Kir6.2 conferred resistance of the KATP channel to SpTx1 inhibition (Figure 3H). Notably, only *hs*Kir6.2 but not Kir6.2 from other species has glutamate at 108 posistion (Figure S4N). These findings confirm the crucial roles of these polar interactions and are largely consistent with previous mutagenesis results ^14,15^.

**Figure 3.**
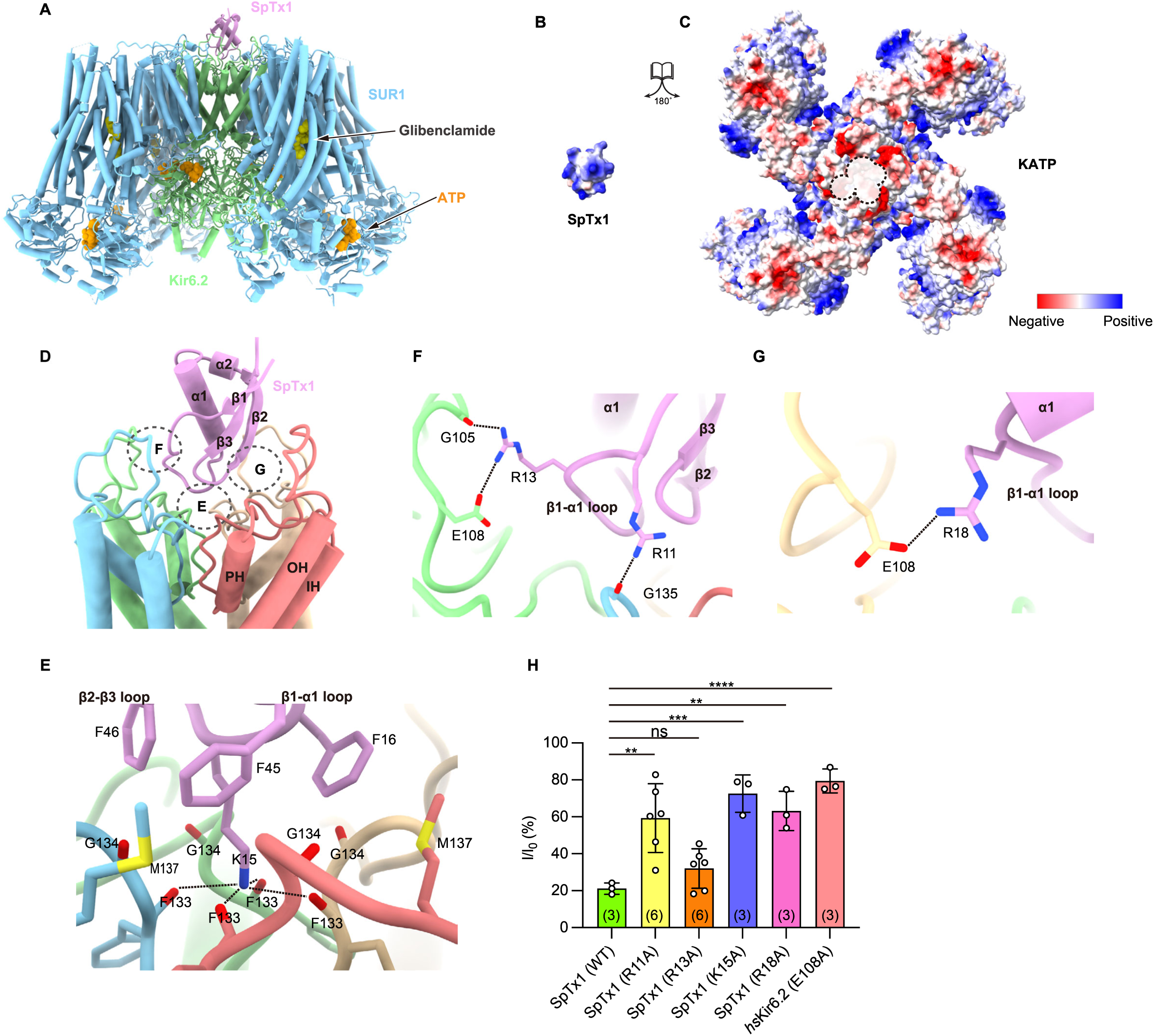
Interaction interface between SpTx1 and KATP. (A) Overall structure of KATP in complex with SpTx1, α-helices, and β-sheets are represented as cylinders and arrows, respectively. The ligands are shown as spheres. The *hs*Kir6.2 subunits, *ma*SUR1 subunits, ATP, Glibenclamide, and SpTx1 are colored in green, blue, orange, yellow, and pink, respectively. (B-C) The open-book view of the interface between KATP (C) and SpTx1 (B) shows the electrostatic potential distribution. The binding sites of SpTx1 to KATP are circled with a dashed line and denoted by a semi-transparent white background. Negatively and positively charged patches are shown in red and blue. (D) SpTx1 and the four *hs*Kir6.2 subunits are shown as cartoons. Key interfaces are circled with dashed lines. SpTx1 and the *hs*Kir6.2 subunits are colored in pink, blue, green, coral, and brown, respectively. IH, inner helix. PH, pore helix. OH, outer helix. (E-G) Detailed views of the interfaces between SpTx1 and *hs*Kir6.2, as circled in (D). (H) Residual currents of *hs*Kir6.2-containing KATP channel inhibited by SpTx1 and its mutants at 1 µM concentration, determined using whole-cell patch clamp. Currents after the SpTx1 application are normalized to the currents before application. Data are presented as mean values ± SD. The numbers of independent experiments are labelled in brackets. *P*-values for inhibition between SpTx1(WT) and SpTx1(R11A), SpTx1(R13A), SpTx1(K15A), and SpTx1(R18A) for the *hs*Kir6.2-containing KATP channel are 1.80 × 10⁻³, 5.99 × 10⁻^1^, 4.00 × 10⁻^4^, and 2.70× 10⁻^3^, respectively. The *P*-value for inhibition between SpTx1(WT) for the *hs*Kir6.2-containing KATP channel and the *hs*Kir6.2(E108A)-containing KATP channel is lower than 1.0 × 10⁻^4^ (\**P* < 0.05, \*\**P* < 0.01, \*\*\**P* < 0.001, \*\*\*\**P* < 0.0001, one-way ANOVA).

### SpTx1 inhibits *hs*Kir6.2 by blocking its ion permeation pathway

KATP channel conducts potassium currents by providing an ion permeation pathway through its pore formed by four Kir6.2 subunits (Figure 4A). The structure of the Kir6.2 pore is highly similar to that of other potassium channels, exemplified by the KcsA ^21^. In the high-resolution crystal structure of KcsA in high potassium condition, potassium ions bind at both the extracellular mouth and four positions (S1-S4) within the selectivity filter ^21^ (Figure 4B).

**Figure 4.**
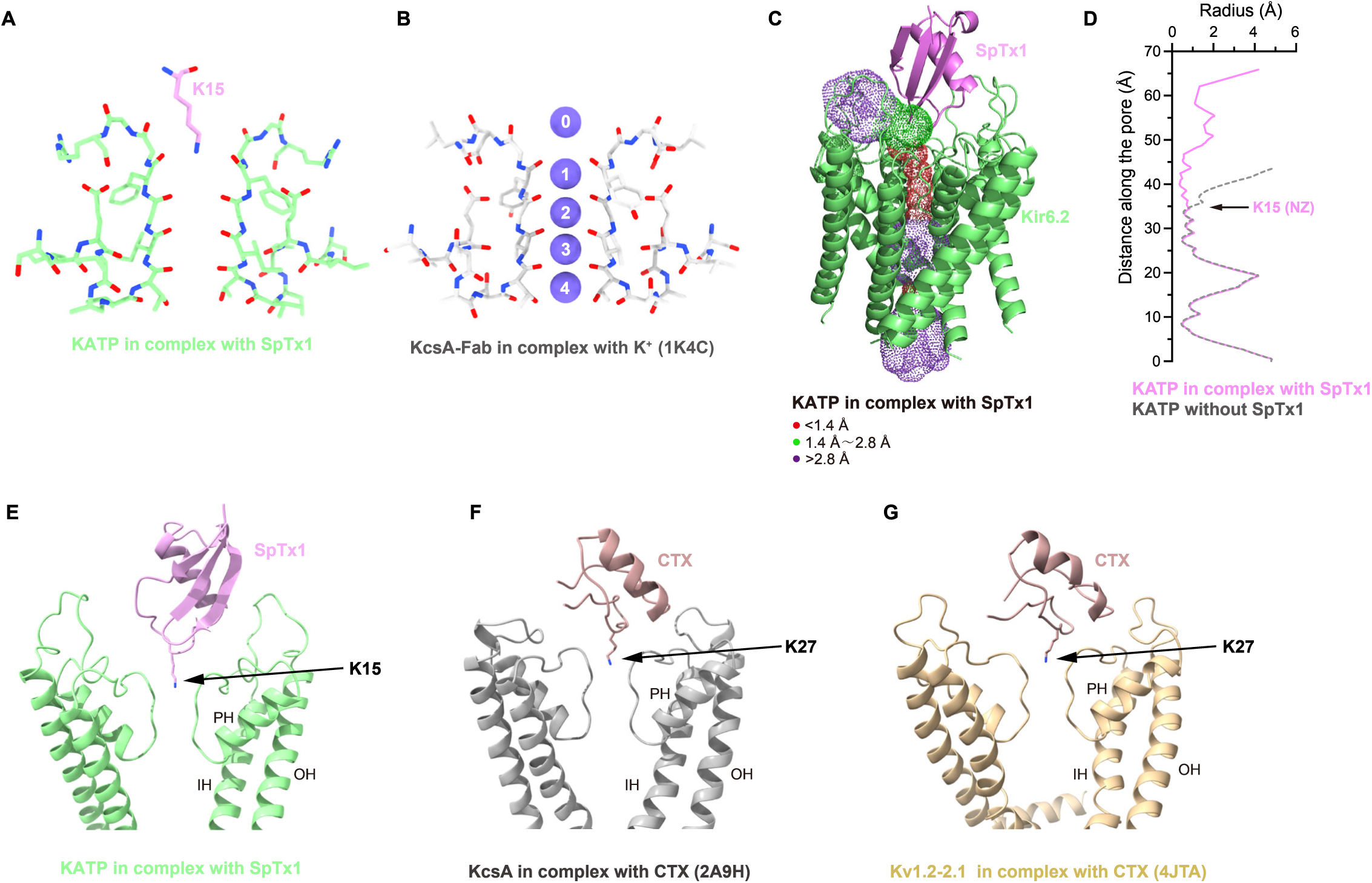
SpTx1 blocks the ion permeation pathway of *hs*Kir6.2. (A) The relative position of K15 of SpTx1 (pink) and the selectivity filter of *hs*Kir6.2 (green). Only two diagonal subunits of *hs*Kir6.2 are shown for clarity. (B) Potassium ion binding sites of the KcsA channel. KcsA and potassium ions are colored in gray and purple. Numbers 0 - 4 denote the ion binding sites along the selectivity filter. (C) The ion permeation pathway along the pore of *hs*Kir6.2 is shown as dots, which are colored in red, green, and purple according to the radii of <1.4, 1.4–2.8, and >2.8 Å. SpTx1 and *hs*Kir6.2 are shown in pink and green cartoons. (D) Pore profiles of *hs*Kir6.2 with (pink line) or without (gray dashed line) SpTx1. The position of the NZ atom of SpTx1-K15 is indicated by an arrow. (E-G) The similar block mechanism of CTX and SpTx1 towards potassium channels. KATP, KcsA, Kv1.2-2.1, SpTx1, and CTX are colored in green, gray, gold, pink, and brown, respectively. The key residues of the toxin are denoted by arrows.

However, in the structure of KATP in complex with SpTx1, SpTx1 physically occupies not only the potassium binding site at the extracellular mouth (P0 site) but also the P1 position in the selectivity filter, precluding the binding of potassium ions at these sites (Figure 4A). Moreover, SpTx1 also occludes the ion permeation pathway of *hs*Kir6.2 (Figure 4A-D), preventing ion permeation (Figure 4D). In the structure, the gate of *hs*Kir6.2 is tightly closed (Figure 4D), in agreement with the binding of inhibitory ATP (Figure 2D). We also found that there is no conformational change in the extracellular regions of Kir6.2 channel upon the binding of SpTx1 (Figure S4K-M). Collectively, these findings indicate that SpTx1 interacts with *hs*Kir6.2 in both its open and closed conformations.

It is previously found that the potassium channel blocker charybdotoxin (CTX), derived from scorpions, also exhibits the α/β defensin fold, characterized by three disulfide bonds. But the structure of CTX is largely different from SpTx1, especially the position of the helix in relation to the rest of the protein (Figure S1F). NMR studies ^22^ or crystallographic studies ^23^ have shown that CTX also binds to the extracellular mouth of the potassium channel pore, with a lysine side chain obstructing the ion permeation pathway (Figure 4F-G), using a mechanism similar to the SpTx1 observed here. However, the orientation of CTX and detailed interactions with potassium are largely different compared to SpTx1 with Kir6.2 (Figure 4E-G).

### Centipede toxin Sm3a is a novel KATP channel blocker

Based on our structural and functional analysis (Figure 3), we identified key residues within the SpTx1 toxin that are essential for inhibiting human Kir6.2 (Figure 1E). Using the SpTx1 sequence as a query, we performed a BLASTp search against the NCBI ClusteredNR database. Among the resulting hits, the toxin Sm3a from the centipede *Scolopendra morsitans* ranked top. Although Sm3a shares only 56% sequence identity with SpTx1, all key residues critical for Kir6.2 inhibition are conserved in Sm3a (Figure 5A). To assess whether Sm3a also inhibits KATP channels, we recombinantly expressed and purified the toxin. Whole-cell patch-clamp recordings demonstrated that Sm3a strongly blocks currents from KATP channels composed of *hs*Kir6.2 and *ma*SUR1 subunits (Figure 5B). Furthermore, application of 200 nM Sm3a inhibited KATP channels slightly better than 200 nM SpTx1 (Figure 5C), indicating comparable inhibitory potency. Given the significant sequence similarity between SpTx1 and Sm3a, we propose that Sm3a binds to and inhibits hsKir6.2-containing KATP channels in a manner similar to SpTx1.

**Figure 5.**
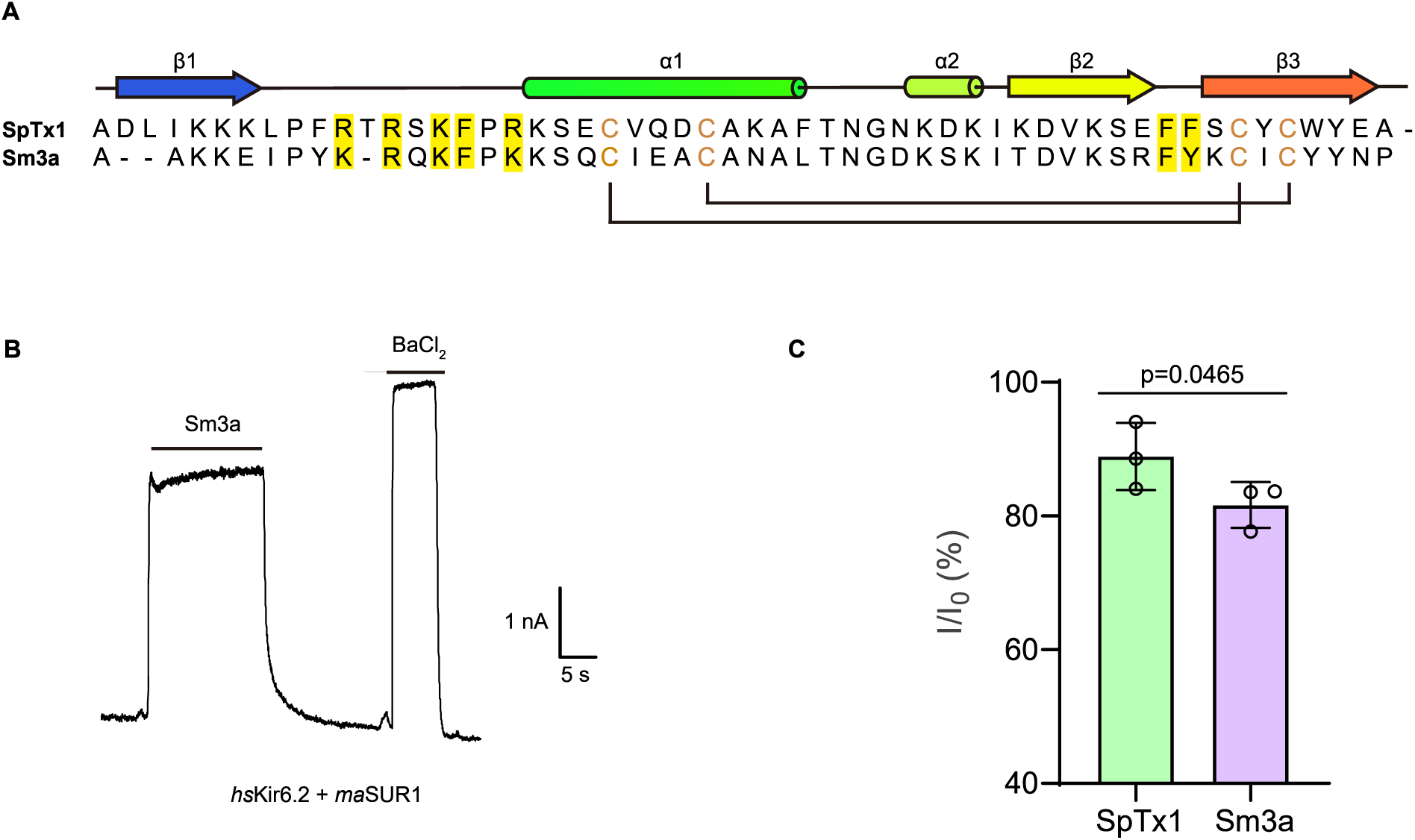
Sm3a inhibits KATP channel. (A) Sequence alignment of SpTx1 and Sm3a. Secondary structures are shown the same as Figure 1E. Residues that are important for KATP inhibition are highlighted in yellow. (B) 1 µM Sm3a inhibits the whole-cell current of the *hs*Kir6.2+*ma*SUR1 KATP channel. Cells were clamped at -40 mV. 5mM BaCl_2_ was used to determine the baseline. (C) Residual currents of *hs*Kir6.2+*ma*SUR1 KATP channel inhibition by 200 nM SpTx1 and Sm3a respectively, determined using whole-cell patch clamp. Currents after SpTx1 and Sm3a application are normalized to the currents before application. Data are presented as mean values ± SD. P-value was calculated with paired t-test.

## Discussion

The structure of SpTx1 determined here reveals a characteristic two-disulfide bond-stabilized α/β defensin fold, which is commonly found in centipede toxins^24^. This structural motif is notable for its stability and functionality, allowing these peptides to interact effectively with various biological targets^17^. The elucidation of complex structure between SpTx1 and *hs*Kir6.2-containing KATP channels provides critical insights into the molecular mechanisms underpinning the inhibitory action of SpTx1. It is known that several gain-of-functional mutations associated with neonatal diabetes, located in either Kir6.2 or SUR1, confer resistance to inhibition by ATP or sulfonylureas (Figure 6A-B) ^5^. The binding site of SpTx1, however, is on the extracellular side of *hs*Kir6.2, distal from the common neonatal diabetes-associated mutations (Figure 6C). This distinct binding mode explains SpTx1’s broad inhibitory effect across diverse clinical KATP channel mutants.

**Figure 6.**
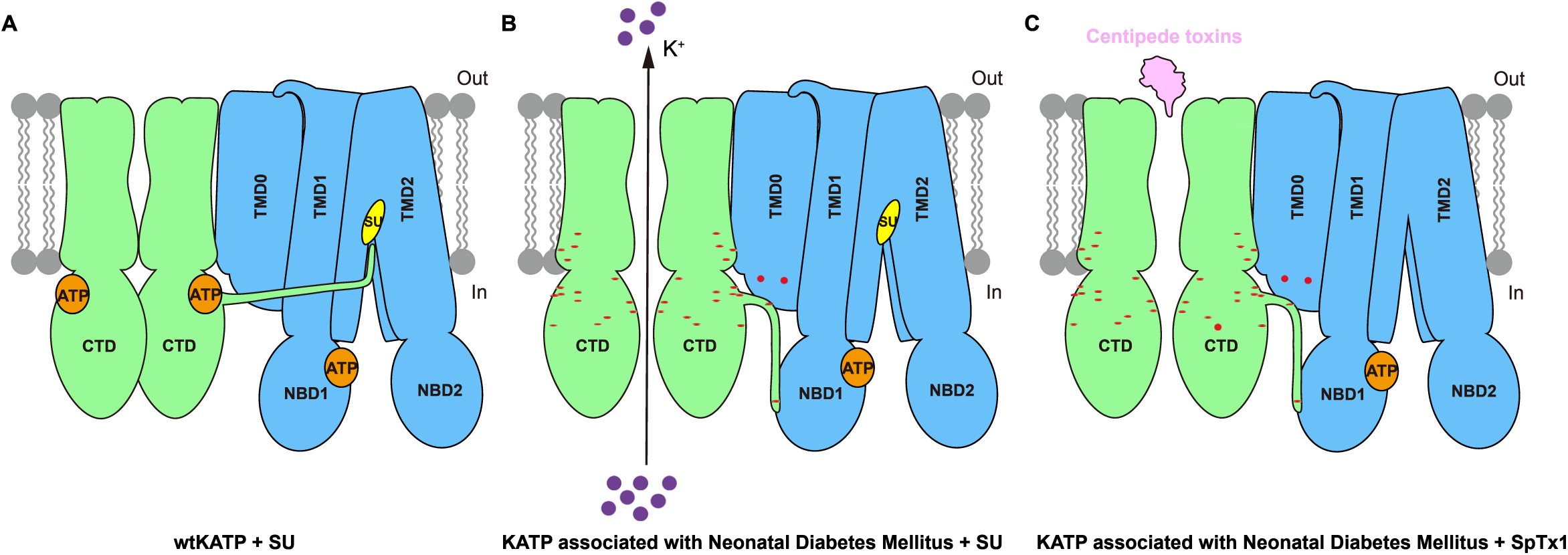
Inhibition mechanism of centipede toxins to KATP channel. (A) Wild-type KATP channels can be effectively inhibited by ATP and sulfonylurea (SU). ATP, SU, Kir6.2, and SUR1 are colored in orange, yellow, green, and blue, respectively. Only two subunits of Kir6.2 and one subunit of SUR1 are shown for clarity. SU recruits KNtp to allosterically close the pore of Kir6.2. SU also inhibits the closure of SUR1 subunits. (B) Some neonatal diabetes mutants of the KATP channel are constitutively active and resistant to ATP and sulfonylurea inhibition. The positions of mutations are indicated with red dots. (C) SpTx1 or Sm3a can effectively block the permeation pathway of KATP channels, including those sulfonylurea-resistant neonatal diabetes mutants.

Moreover, the molecular targets of defensin fold centipede toxins are diverse, many of which target different potassium channels^25^. The structure mechanism of interactions between Kir6.2 and SpTx1 could serve as a paradigm for investigating other defensin fold toxin-channel interactions, potentially leading to the discovery of novel inhibitors or modulators for clinical applications. Indeed, structure-based sequence search aids the identification of Sm3a as a new inhibitor of KATP channel (Figure 5).

Importantly, SpTx1 and KATP interact at the extracellular side of the Kir6.2 subunit and neither the N-nor the C-termini of SpTx1 is involved in its binding to Kir6.2. As a result, there is a unique opportunity to explore protein fusions to either the N- or C-terminus of SpTx1 without compromising its functionality. This opens avenues for new modifications of SpTx1, such as fusing SpTx1 with proteins that target endocytosis receptors to facilitate the selective degradation of gain-of-function KATP mutants ^26,27^. Targeting these mutants through the lysosomal degradation pathway might represent a novel therapeutic strategy, leveraging the specificity of SpTx1’s binding properties.

In conclusion, the structural and functional insights gained from the study of SpTx1-KATP complex not only enhance our understanding of the mechanism of centipede toxins but also lay the groundwork for potential therapeutic modifications of these centipede toxins to enhance their potency or broaden its application scope.

## Methods

### Constructs

CDS of the matured form of SpTx1 and Sm3a (P0DQC2.1) were synthesized and inserted into a pET-based N-terminally His-tagged vector (pPH) with TEV protease cleavage site to generate the toxin expression vectors.

CDS of maSUR1 (E203K) and hsKir6.2 (Q52E) are linked by a long linker including a PreScission protease cleavage site (VDGSGSGSGSAAGSGSGSGSGSGAAGSGSGSGSGSGAAALELEVLFQGP). The initial Methionine of *hs*Kir6.2 after the linker was removed. The ORF of fusion protein was inserted into a modified BacMam vector with C-terminally GFP and two Strep tags^28^.

### Expression and purification of toxin

The expression vector for the toxins were transformed into Rosetta-gami 2(DE3) competent cells. Protein expression was induced by adding 1 mM IPTG for 3 hours at 37°C in LB medium. Collected bacterial cells were resuspended in 20 mM Tris (pH 7.5), 150 mM NaCl, 1 mM PMSF and were disrupted via sonication. The lysate was centrifuged at 30,000g for 1.5 hours, after which the supernatant was applied to a Talon affinity column and eluted using 250 mM imidazole. The eluted protein was subjected to TEV protease digestion to remove the purification tag. After cleavage, contaminant protein was precipitated using methanol, yielding a highly purified toxin protein, which was subsequently purified via SP ion-exchange chromatography to remove residual methanol. Final purification was performed using a Superdex 75 size-exclusion column running in 10 mM Tris (pH 7.5) and 150 mM NaCl. Protein was concentrated to 20 mg/ml for crystallization.

### Crystallization, X-ray data collection and structural determination

Protein was mixed with 0.1 M succinic acid pH 5.5, 0.05 M ammonium sulfate, 30% (v/v) pentaerythritol ethoxylate (15/4 EO/OH) with 1/1 ratio at room temperature in a hanging drop crystallization setup. Crystal was harvested using 0.1 M succinic acid pH 5.5, 0.05 M ammonium sulfate, 30% (v/v) pentaerythritol ethoxylate (15/4 EO/OH), 10 % ethylene glycol as cryoprotectant and flash frozen in liquid nitrogen.

Native dataset of SpTx1 were collected at Shanghai Synchrotron Radiation Facility beamline BL17U at 0.97918 Å. Diffraction images were processed using HKL3000^29^ and converted to mtz file using CCP4^30^. SpTx1 structure predicted by AlphaFold2 ^31^ was used as the search model for molecular replacement using Phaser ^32^. The model was rebuilt manually using COOT ^33^ and was refined using Phenix ^34^. The final model contains 99.68 % favored and 0.32 % allowed residues but no outliers on the Ramachandran plot.

### Expression and purification of KATP channel

KATP channels were expressed as described previously and the purification process was carried out with minor modification^7,35^. For protein purification, membrane pellets were homogenized in TBS (20 mM Tris-HCl pH 7.5, 150 mM NaCl) and solubilized in TBS with 1% digitonin (Calbiochem), supplemented with protease inhibitors (1 mg/ml Leupeptin, 1 mg/ml Pepstatin, 1 mg/ml Aprotinin, and 1 mM PMSF), 5 μM glibenclamide, 1 mM ATP, 1 mM EDTA, 2.5 mM MgCl₂, 2.5 mM CaCl₂, for 30 min at 4℃. Insoluble materials were removed after centrifugation at 100,000 g for 30 min and the supernatant was loaded onto a 5 mL column packed with Streptactin 4FF resin (Smart Lifesciences). Strep column was first washed by buffer A (5 μM GBM, 1 mM ATP, 1 mM EDTA, 0.1% digitonin in TBS), then 1 mM ATP and 2.5 mM MgCl₂ in buffer A and finally in buffer A again. KATP protein was eluted with 10 mM desthiobiotin in buffer A. Protein was then digested with PreScission protease overnight, concentrated, and purified using Superose 6 column running in buffer A. Peak fractions were collected and concentrated to A_280_=15 (estimated as 15 µM K_ATP_ octamers) for cryo-EM sample preparation.

### Cryo-EM sample preparation

KATP octamers were supplemented with 200 µM GBM and 1 mM ATP, along with 100 µM SpTx1. The final KATP protein concentration was estimated to be 9 µM octamer in solution. Cryo-EM grids were prepared with GIG R1/1 holey carbon grids, which were glow-discharged for 120 s. 2.5 µl KATP octamers sample was applied to the glow-discharged grid and then the grid was blotted at blotting force in level 2 for 2 s at 100% humidity and 20 °C, before plunge-frozen into the liquid ethane.

### Cryo-EM data collection

Cryo-EM data collection was performed on a Titan Krios microscope (Thermo Fisher Scientific) operating at 300 kV. Images were acquired using a Gatan K2 direct electron detector mounted post a Quantum energy filter with a 20 eV slit width. Super-resolution mode was employed with a calibrated pixel size of 1.324 Å at the specimen level, and data acquisition was controlled via Serial EM software. Defocus values were systematically varied between −1.3 μm and −1.8 μm to optimize contrast transfer. A dose rate of 8 e⁻/s/Å² was applied, with a cumulative electron exposure of 50 e⁻/Å². Each 12-second exposure was dose-fractionated into 50 frames to facilitate motion correction during image processing.

### Cryo-EM image processing and model building

5,655 movies were corrected for beam-induced motion, dose-weighted, and two-fold binned to a pixel size of 1.324 Å using MotionCor2 ^36^. The contrast transfer function (CTF) parameters of micrographs were estimated using patch CTF function in cryoSPARC ^37^. Iterative 2D and 3D classification generated 548,390 particles with 256 box size which were refined to 2.74 Å overall resolution using non-uniform refinement function in cryoSPARC^37^ . Symmetry expansion, particle subtraction and local refinement of the SUR1_ABC_ regions yielded a local map of 2.87 Å resolution. Symmetry expansion and iterative focused classification of SpTx1 region using C1 symmetry resolved two classes: one has SpTx1 bound and the other has no SpTx1 bound. The class with SpTx1 bound was locally refined using C1 symmetry focusing on the KATP_CORE_ region to obtain a map at 2.88 Å resolution. This map of KATP_CORE_ was combined with the map of SUR1_ABC_ to obtain a composite map for model building and interpretation. The structures of SpTx1 and KATP channel in complex with repaglinide (PDB ID: 6JB1) were docked into the composite map. The model was rebuilt manually using COOT ^33^ and was refined using Phenix^34^.

### Electrophysiology

KATP constructs were transfected into FreeStyle 293-F cells using polyethylenimine at a cell density of 1 × 10^6^ cells/ml. Cells were cultured in FreeStyle 293 Expression Medium with 1% FBS for 24h before recording. Currents were recorded using Whole-cell mode at -40 mV in the pipette and an Axon-patch 200B amplifier (Axon Instruments, USA). Patch electrodes were pulled by a horizontal micro-electrode puller (P-1000, Sutter Instrument Co, USA) to tip resistance of 2.0–3.0 MΩ. Pipette solution containing (mM): 107 KCl, 1.2 MgCl_2_, 1 CaCl_2_, 10 EGTA, 5 HEPES (pH 7.2, KOH) and bath solution containing (mM): 40 KCl, 100 NaCl, 2.6 CaCl_2_, 1.2 MgCl_2_, 5 HEPES (pH 7.4, NaOH) were used for measuring inhibitory effect of wild-type and mutant toxins. Application of 5 mM BaCl_2_ in bath solution was used to block the KATP currents to determine the baseline. Data analysis was carried out using GraphPad Prism 8.0 or 10.4.0.

### Quantification and statistical analysis

Global resolution estimations of cryo-EM density maps are based on the 0.143 Fourier Shell Correlation criterion ^38^. The local resolution was estimated using cryoSPARC. The number of independent experiments (N) and the relevant statistical parameters for each experiment (such as mean or standard deviation) are described in the figure legends. No statistical methods were used to pre-determine sample sizes.

## Data Availability

Crystal structure of SpTx1 has been deposited into PDB under ID codes: 9KGA. Cryo-EM maps and the atomic coordinate of the SpTx-KATP complex have been deposited in the EMDB and PDB under the ID codes EMDB: EMD-62322 and PDB ID: 9KGL.

## Acknowledgments

We thank all of the Chen Lab members for their kindly help. Cryo-EM data collection was supported by the Electron microscopy laboratory, and the Cryo-EM platform of Peking University. Part of the structural computation was also performed on the Computing Platform of the Center for Life Science and High-performance Computing Platform of Peking University. We thank the National Center for Protein Sciences at Peking University in Beijing, China for assistance with negative stain EM. We thank staffs at Shanghai Synchrotron Radiation Facility beamline BL17U for help with X-ray crystal diffraction data. The work is supported by grants from the Ministry of Science and Technology of China (National Key R&D Program of China, 2022YFA0806504 to L.C.), National Natural Science Foundation of China (31570762 to C.F. and 32225027 to L.C.), and the Center For Life Sciences (CLS to L.C.).

## Author contributions

L.C. initiated the project. M.W. purified the SpTx1 protein, carried out crystallization, prepared the cryo-EM sample, collected the cryo-EM data. M.W., T.H., and L.C. collected the X-ray diffraction data. C.F. and L.C. determined the crystal structure of SpTx1. L.C. processed the cryo-EM data and built the model. T.H. and Y. T. purified centipede toxins and their mutants and carried out electrophysiological experiments. All authors contributed to the manuscript preparation.

---

**Figure S1.**
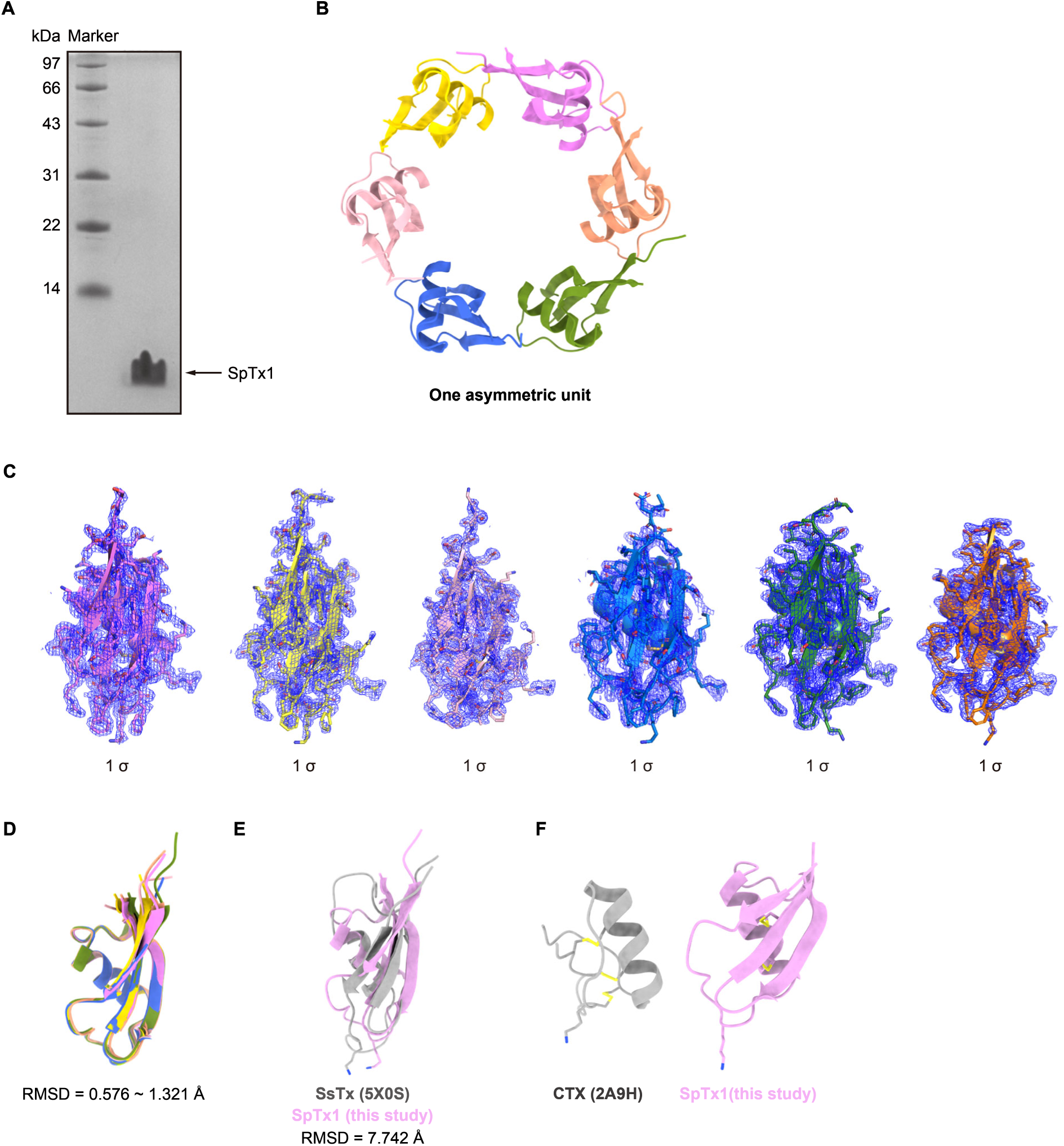
Crystal structure of SpTx1. (A) Coomassie blue-stained SDS-PAGE of SpTx1 used for crystallization. (B) One asymmetric unit of SpTx1 is shown as a hexametric ring. (C) Density of each monomer SpTx1 in (B). (D) Aligned monomers with root mean square deviation 0.576-1.321Å. (E) Structural comparison between SpTx1 and SsTx with root mean square deviation 7.742Å, SsTx and SpTx are colored in gray and pink, respectively. (F) CTX and SpTx1 are colored in gray and pink, respectively. Disulfide bonds are shown as sticks.

**Figure S2.**
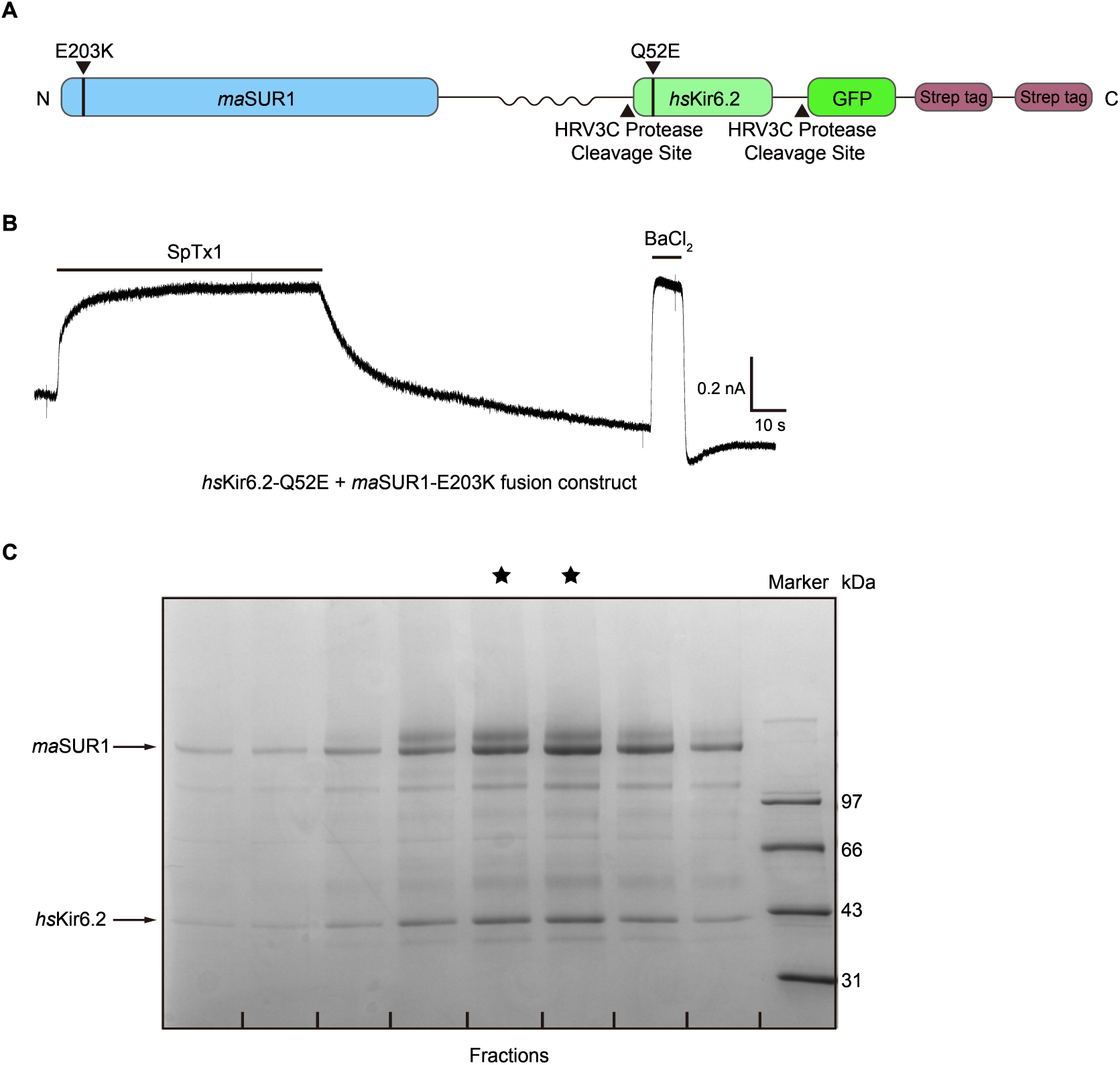
Protein expression and purification of KATP. (A) The design of the KATP fusion construct used for Cryo-EM. *ma*SUR1, *hs*Kir6.2, GFP, and Strep-tag are colored in blue, pale green, green, and brown, respectively. A black wavy line represents the linker between *ma*SUR1 and *hs*Kir6.2. PreScission Protease cleavage sites and mutation sites are indicated by black triangles. (B) Inhibition of the KATP fusion channel by SpTx1, determined using whole-cell patch clamp. (C) Coomassie blue-stained SDS-PAGE of the KATP protein sample from size-exclusion chromatography. Fractions labeled with asterisks were pooled for cryo-EM sample preparation.

**Figure S3.**
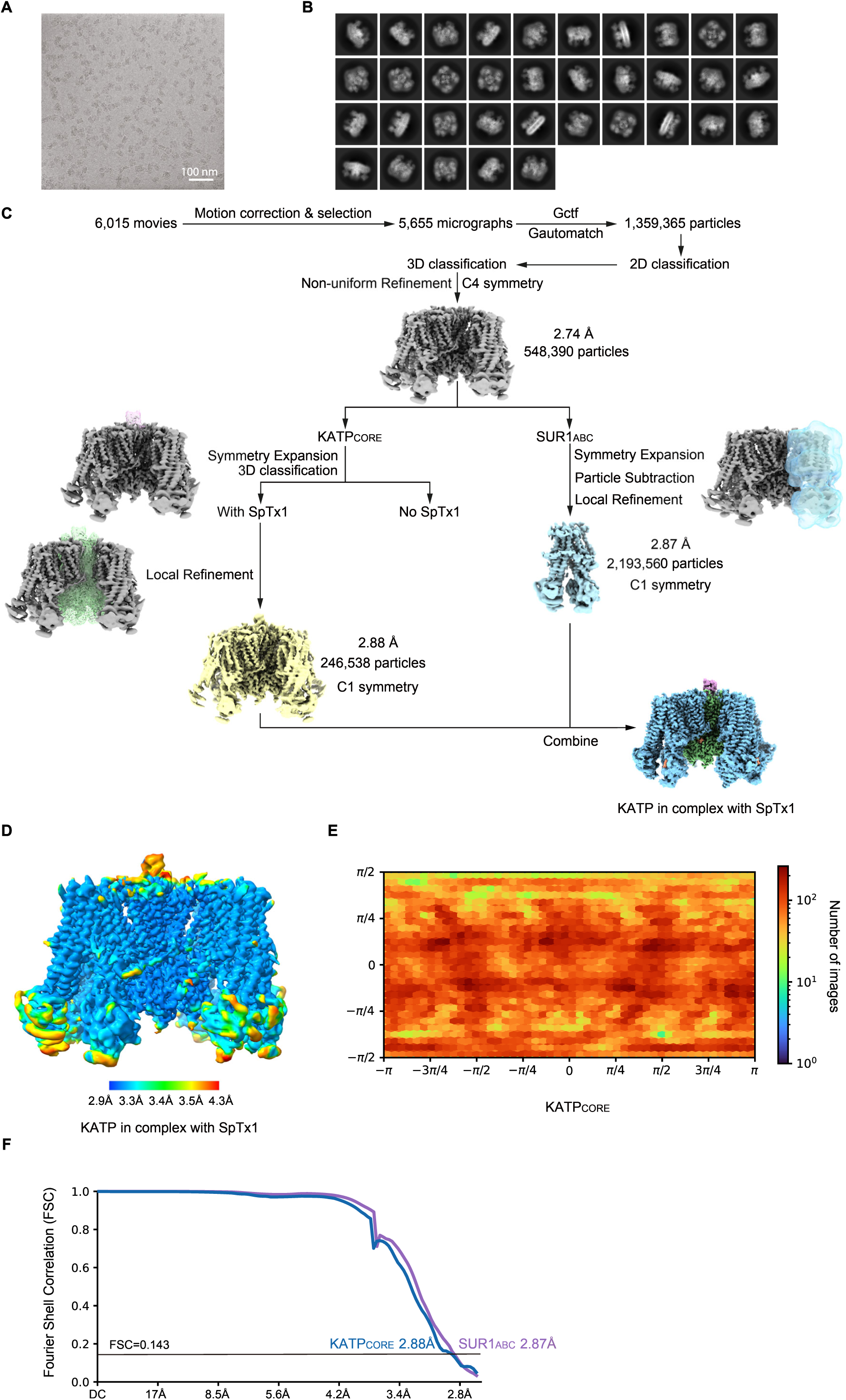
Workflow for cryo-EM data processing of KATP in complex with SpTx1. (A) Representative image from a dataset consisting of 6,015 motion-corrected micrographs. (B) Two-dimensional class averages. (C) Cryo-EM image processing workflow. (D) Local resolution map of the composite map. (E) Angular distribution of the KATP_CORE_. (F) Resolution estimation of the KATP_CORE_ and SUR1_ABC_ map, based on the gold-standard FSC 0.143 cut-off criterion.

**Figure S4.**
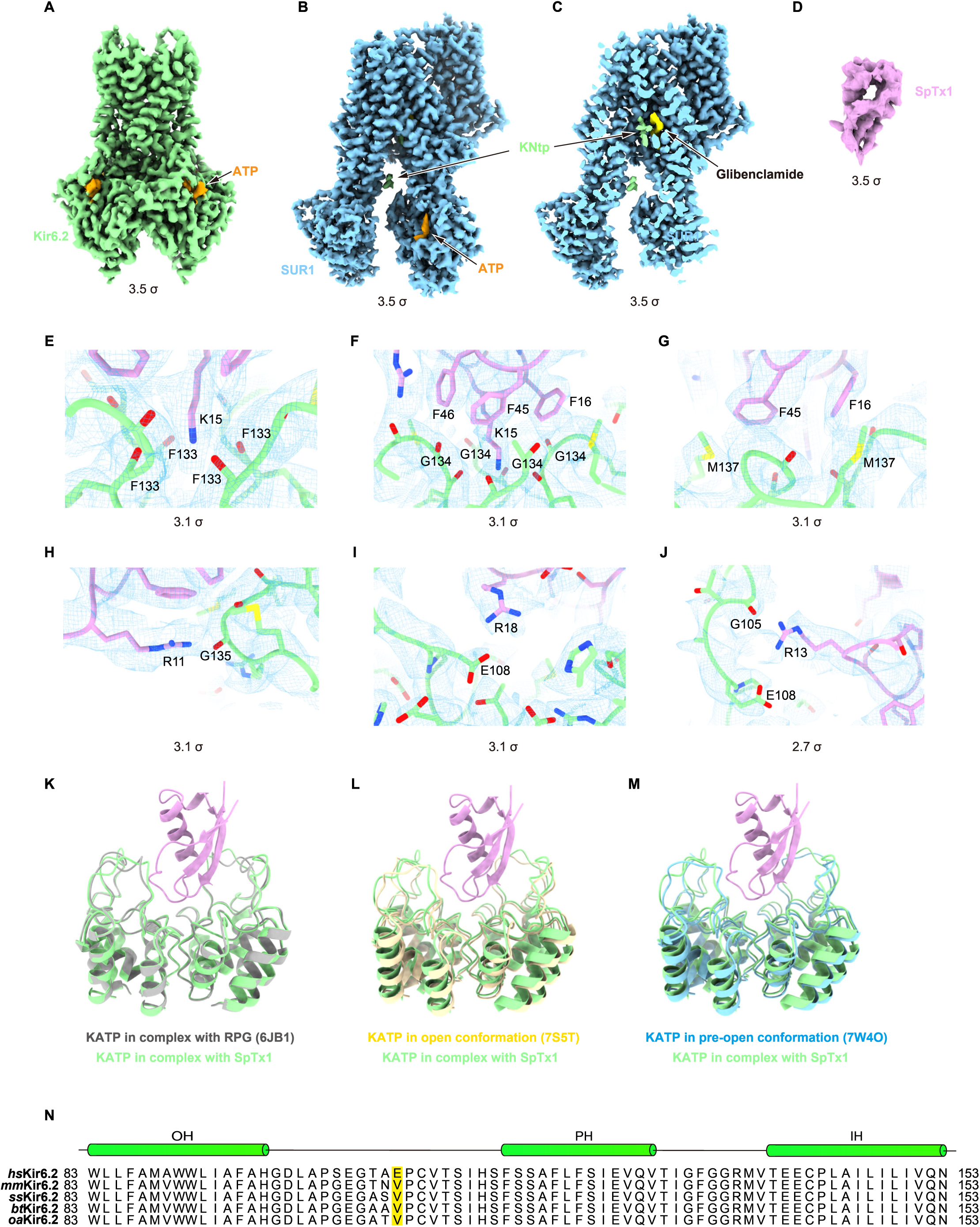
Local cryo-EM densities of KATP in complex with SpTx1. (A) Density map of the *hs*Kir6.2 subunit, contoured at 3.5 σ, viewed from the side. (B) Density map of the *ma*SUR1 subunit, contoured at 3.5 σ, viewed from the side. (C) Cut-open view of (B). (D) Density map of SpTx1, contoured at 3.5 σ, viewed from the side. (E-J) Close-up views of the key interfaces between SpTx1 and *hs*Kir6.2. SpTx1 is colored in pink, *hs*Kir6.2 is colored in green, and the density map is shown as a blue mesh, contoured at 3.1 σ and 2.7 σ. (K-M) Structure of KATP in complex with SpTx1 compared with the SpTx1-free closed state (K), the open state (L), and the pre-open state (M) KATP, colored in green, gray, yellow, and blue, respectively. (N) Sequence alignment of Kir6.2 from Homo sapiens (*hs*Kir6.2), Mus musculus (*mm*Kir6.2), Susscrofa (*ss*Kir6.2), Bos taurus (*bt*Kir6.2), and Ovis aries (*oa*Kir6.2). α-helices are represented as green cylinders. Key residues are indicated by a yellow background.

**Figure S5.**
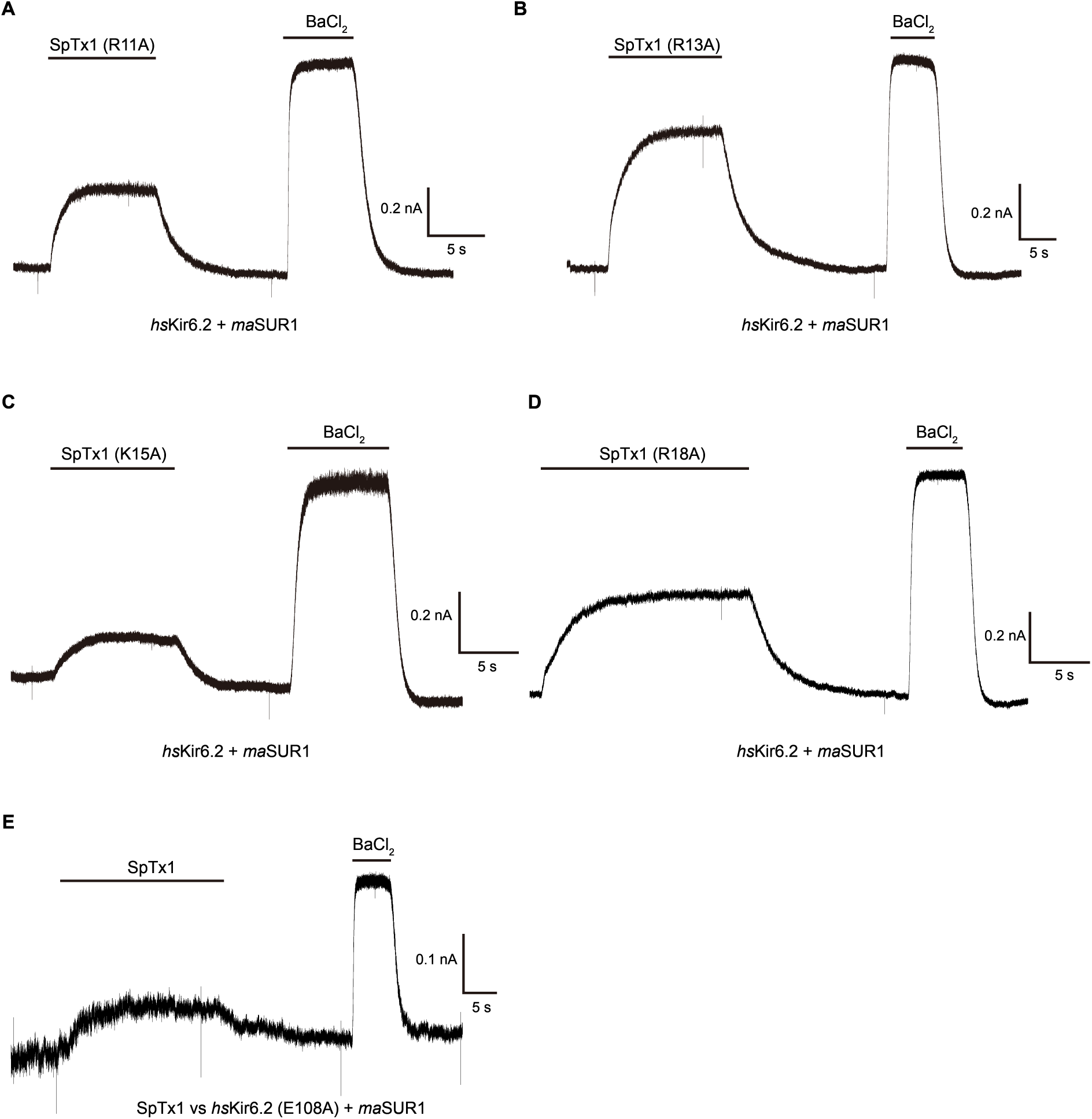
Representative whole-cell currents of KATP inhibition by various SpTx1 mutants. (A) Inhibition of the wild-type hsKir6.2+maSUR1 KATP channel by the SpTx1 R11A mutant. (B) Inhibition of the wild-type hsKir6.2+maSUR1 KATP channel by the SpTx1 R13A mutant. (C) Inhibition of the wild-type hsKir6.2+maSUR1 KATP channel by the SpTx1 K15A mutant. (D) Inhibition of the wild-type hsKir6.2+maSUR1 KATP channel by the SpTx1 R18A mutant. (E) Inhibition of the hsKir6.2 E108A mutant + maSUR1 KATP channel by wild-type SpTx1.

**Table S1.** Data collection and refinement statistics of SpTx1 (molecular replacement)

| SpTx1 (PDB ID: 9KGA) |  |
| --- | --- |
| **Data collection** |  |
| Space group | *P3<sub>1</sub>21* |
| Cell dimensions |  |
| *a*, *b*, *c* (Å) | 60.49, 60.49, 169.85 |
| $\alpha$, $\beta$, $\gamma$ (°) | 90.00, 90.00, 120.00 |
| Resolution (Å) | 50-1.90 (1.93-1.90)* |
| *R*<sub>sym</sub> or *R*<sub>merge</sub> | 0.11 (0.75) |
| *I* / $\sigma I$ | 22.50 (3.33) |
| Completeness (%) | 100.00(100.00) |
| Redundancy | 9.2 (8.8) |
| **Refinement** |  |
| Resolution (Å) | 44.58-1.9 |
| No. reflections | 29,121 |
| *R*<sub>work</sub> / *R*<sub>free</sub> | 0.2031/0.2138 |
| No. atoms | 3,000 |
| Protein | 2,702 |
| Ligand/ion | 0 |
| Water | 298 |
| *B*-factors (Å<sup>2</sup>) | 28.54 |
| Protein | 27.99 |
| Ligand/ion |  |
| Water | 33.51 |
| R.m.s. deviations |  |
| Bond lengths (Å) | 0.01 |
| Bond angles (°) | 0.81 |

\*Values in parentheses are for highest-resolution shell.

**Table S2.** Cryo-EM data collection, refinement and validation statistics.

|  | Composite map<br>(EMD-62322)<br>(PDB 9KGL) | Focused map of<br>KATP<sub>core</sub><br>(EMD-62311) | Focused map of<br>SUR1<sub>ABC</sub><br>(EMD-62317) |
| --- | --- | --- | --- |
| **Data collection and processing** |  |  |  |
| Magnification |  | 105,000 × |  |
| Voltage (kV) |  | 300 |  |
| Electron exposure (e<sup>-</sup>/Å<sup>2</sup>) |  | 50 |  |
| Defocus range (μm) |  | -1.5 to -1.8 |  |
| Pixel size (Å) |  | 1.324 |  |
| Symmetry imposed |  | C1 |  |
| Initial particle images (no.) |  | 1,595,249 |  |
| Final particle images (no.) |  | 246,583 | 2,193,560* |
| Map resolution (Å) |  | 2.88 | 2.87 |
| FSC threshold |  | 0.143 | 0.143 |
| Map resolution range (Å) |  | 250-2.88 | 250-2.87 |
| **Refinement** |  |  |  |
| Initial model used (PDB code) | 6JB1 |  |  |
| Model resolution (Å) | 3.1 |  |  |
| FSC threshold | 0.5 |  |  |
| Model resolution range (Å) | 250-3.1 |  |  |
| Map sharpening *B* factor (Å<sup>2</sup>) |  | 112.4 | 147.9 |
| Model composition |  |  |  |
| Non-hydrogen atoms | 55,831 |  |  |
| Protein residues | 7,074 |  |  |
| Ligands | 12 |  |  |
| *B* factors (Å<sup>2</sup>) |  |  |  |
| Protein | 69.12 |  |  |
| Ligand | 58.30 |  |  |
| R.m.s. deviations |  |  |  |
| Bond lengths (Å) | 0.005 |  |  |
| Bond angles (°) | 0.996 |  |  |
| Validation |  |  |  |
| MolProbity score | 1.65 |  |  |
| Clashscore | 5.66 |  |  |
| Poor rotamers (%) |  |  |  |
| Ramachandran plot |  |  |  |
| Favored (%) | 97.61 |  |  |
| Allowed (%) | 2.27 |  |  |
| Disallowed (%) | 0.11 |  |  |

\* The particles are expended using C4 symmetry.

**Video S1. Cryo-EM structure of KATP channel in complex with SpTx1**

The cryo-EM map and atomic model of KATP and SpTx1 complex is colored the same as Fig. 2a and rotated to show different views.

## References

1 Nichols, C. G. KATP channels as molecular sensors of cellular metabolism. Nature 440, 470–476 (2006). 10.1038/nature04711

2 Sakura, H., Ammala, C., Smith, P. A., Gribble, F. M. & Ashcroft, F. M. Cloning and functional expression of the cDNA encoding a novel ATP-sensitive potassium channel subunit expressed in pancreatic beta-cells, brain, heart and skeletal muscle. FEBS Lett. 377, 338–344 (1995). 10.1016/0014-5793(95)01369-5

3 Schmid, D. et al. An abundant, truncated human sulfonylurea receptor 1 splice variant has prodiabetic properties and impairs sulfonylurea action. Cell. Mol. Life Sci. 69, 129–148 (2012). 10.1007/s00018-011-0739-x

4 Wu, J. X., Ding, D. & Chen, L. The Emerging Structural Pharmacology of ATP-Sensitive Potassium Channels. Mol. Pharmacol. 102, 234–239 (2022). 10.1124/molpharm.122.000570

5 Pipatpolkai, T., Usher, S., Stansfeld, P. J. & Ashcroft, F. M. New insights into KATP channel gene mutations and neonatal diabetes mellitus. Nat Rev Endocrinol 16, 378–393 (2020). 10.1038/s41574-020-0351-y

6 Martin, G. M., Kandasamy, B., DiMaio, F., Yoshioka, C. & Shyng, S. L. Anti-diabetic drug binding site in a mammalian KATP channel revealed by Cryo-EM. Elife 6 (2017). 10.7554/eLife.31054

7 Wu, J. X., Ding, D., Wang, M., Kang, Y., Zeng, X. & Chen, L. Ligand binding and conformational changes of SUR1 subunit in pancreatic ATP-sensitive potassium channels. Protein Cell 9, 553–567 (2018). 10.1007/s13238-018-0530-y

8 Ding, D., Wang, M., Wu, J. X., Kang, Y. & Chen, L. The Structural Basis for the Binding of Repaglinide to the Pancreatic KATP Channel. Cell Rep 27, 1848–1857 e1844 (2019). 10.1016/j.celrep.2019.04.050

9 Martin, G. M. et al. Mechanism of pharmacochaperoning in a mammalian KATP channel revealed by cryo-EM. Elife 8 (2019). 10.7554/eLife.46417

10 Wu, J. X., Ding, D., Wang, M. & Chen, L. Structural Insights into the Inhibitory Mechanism of Insulin Secretagogues on the Pancreatic ATP-Sensitive Potassium Channel. Biochemistry (Mosc.) 59, 18–25 (2020). 10.1021/acs.biochem.9b00692

11 Wang, M., Wu, J. X. & Chen, L. Structural Insights Into the High Selectivity of the Anti-Diabetic Drug Mitiglinide. Front Pharmacol 13, 929684 (2022). 10.3389/fphar.2022.929684

12 Yang, Y. & Chen, L. Functional dissection of KATP channel structures reveals the importance of a conserved interface. Structure (2023). 10.1016/j.str.2023.11.008

13 Ramu, Y., Xu, Y. & Lu, Z. A novel high-affinity inhibitor against the human ATP-sensitive Kir6.2 channel. J. Gen. Physiol. 150, 969–976 (2018). 10.1085/jgp.201812017

14 Ramu, Y., Yamakaze, J., Zhou, Y., Hoshi, T. & Lu, Z. Blocking Kir6.2 channels with SpTx1 potentiates glucose-stimulated insulin secretion from murine pancreatic beta cells and lowers blood glucose in diabetic mice. Elife 11 (2022). 10.7554/eLife.77026

15 Ramu, Y. & Lu, Z. A family of orthologous proteins from centipede venoms inhibit the hKir6.2 channel. Sci Rep 9, 14088 (2019). 10.1038/s41598-019-50688-x

16 Tang, D. et al. Molecular mechanisms of centipede toxin SsTx-4 inhibition of inwardly rectifying potassium channels. J. Biol. Chem. 297, 101076 (2021). 10.1016/j.jbc.2021.101076

17 Luo, L. et al. Centipedes subdue giant prey by blocking KCNQ channels. Proc. Natl. Acad. Sci. U. S. A. 115, 1646–1651 (2018). 10.1073/pnas.1714760115

18 Pratt, E. B., Zhou, Q., Gay, J. W. & Shyng, S. L. Engineered interaction between SUR1 and Kir6.2 that enhances ATP sensitivity in KATP channels. J. Gen. Physiol. 140, 175–187 (2012). 10.1085/jgp.201210803

19 Lee, K. P. K., Chen, J. & MacKinnon, R. Molecular structure of human KATP in complex with ATP and ADP. Elife 6 (2017). 10.7554/eLife.32481

20 Zhao, C. & MacKinnon, R. Molecular structure of an open human KATP channel. Proc. Natl. Acad. Sci. U. S. A. 118 (2021). 10.1073/pnas.2112267118

21 Zhou, Y., Morais-Cabral, J. H., Kaufman, A. & MacKinnon, R. Chemistry of ion coordination and hydration revealed by a K+ channel-Fab complex at 2.0 A resolution. Nature 414, 43–48 (2001). 10.1038/35102009

22 Yu, L. et al. Nuclear magnetic resonance structural studies of a potassium channel-charybdotoxin complex. Biochemistry (Mosc.) 44, 15834–15841 (2005). 10.1021/bi051656d

23 Banerjee, A., Lee, A., Campbell, E. & Mackinnon, R. Structure of a pore-blocking toxin in complex with a eukaryotic voltage-dependent K(+) channel. Elife 2, e00594 (2013). 10.7554/eLife.00594

24 Dash, T. S. et al. A Centipede Toxin Family Defines an Ancient Class of CSalphabeta Defensins. Structure 27, 315–326 e317 (2019). 10.1016/j.str.2018.10.022

25 Du, C. et al. Centipede KCNQ Inhibitor SsTx Also Targets K(V)1.3. Toxins (Basel) 11 (2019). 10.3390/toxins11020076

26 Huang, B. et al. Designed endocytosis-inducing proteins degrade targets and amplify signals. Nature (2024). 10.1038/s41586-024-07948-2

27 Zhang, D. et al. Transferrin receptor targeting chimeras for membrane protein degradation. Nature (2024). 10.1038/s41586-024-07947-3

28 Li, N., Wu, J. X., Ding, D., Cheng, J., Gao, N. & Chen, L. Structure of a Pancreatic ATP-Sensitive Potassium Channel. Cell 168, 101–110 e110 (2017). 10.1016/j.cell.2016.12.028

29 Otwinowski, Z. & Minor, W. Processing of X-ray diffraction data collected in oscillation mode. Meth. Enzymol. 276, 307–326 (1997).

30 Agirre, J. et al. The CCP4 suite: integrative software for macromolecular crystallography. Acta Crystallogr D Struct Biol 79, 449–461 (2023). 10.1107/S2059798323003595

31 Jumper, J. et al. Highly accurate protein structure prediction with AlphaFold. Nature 596, 583–589 (2021). 10.1038/s41586-021-03819-2

32 McCoy, A. J., Grosse-Kunstleve, R. W., Adams, P. D., Winn, M. D., Storoni, L. C. & Read, R. J. Phaser crystallographic software. J Appl Crystallogr 40, 658–674 (2007). 10.1107/S0021889807021206

33 Emsley, P., Lohkamp, B., Scott, W. G. & Cowtan, K. Features and development of Coot. Acta Crystallogr. D Biol. Crystallogr. 66, 486–501 (2010). 10.1107/S0907444910007493

34 Adams, P. D. et al. The Phenix software for automated determination of macromolecular structures. Methods 55, 94–106 (2011). 10.1016/j.ymeth.2011.07.005

35 Wang, M., Wu, J. X., Ding, D. & Chen, L. Structural insights into the mechanism of pancreatic KATP channel regulation by nucleotides. Nat Commun 13, 2770 (2022). 10.1038/s41467-022-30430-4

36 Zheng, S. Q., Palovcak, E., Armache, J. P., Verba, K. A., Cheng, Y. & Agard, D. A. MotionCor2: anisotropic correction of beam-induced motion for improved cryo-electron microscopy. Nat. Methods 14, 331–332 (2017). 10.1038/nmeth.4193

37 Punjani, A., Rubinstein, J. L., Fleet, D. J. & Brubaker, M. A. cryoSPARC: algorithms for rapid unsupervised cryo-EM structure determination. Nat. Methods 14, 290–296 (2017). 10.1038/nmeth.4169

38 Chen, S. et al. High-resolution noise substitution to measure overfitting and validate resolution in 3D structure determination by single particle electron cryomicroscopy. Ultramicroscopy 135, 24–35 (2013). 10.1016/j.ultramic.2013.06.004

